# Evaluation of wound healing effects of ginsenoside Rg1 and red ginseng extract in STZ-induced diabetic wound model: an in vivo pilot study

**DOI:** 10.1101/2021.05.05.442721

**Authors:** Ji Yun Lim, Young Suk Choi, Hye Rim Lee, Hye Min An, Young Koo Lee

**Author notes:** **Corresponding author: Young Koo Lee, MD, PhD** Departments of Orthopedic Surgery, Soonchunhyang University, Bucheon Hospital, Jomaru-ro, Bucheon-si, Gyeonggi-do, Republic of Korea. E-mail address (Y.-K. Lee). Departments of Orthopedic Surgery, Soonchunhyang University, Bucheon Hospital, Jomaru-ro, Bucheon-si, Gyeonggi-do, Republic of Korea, Republic of Korea (Ji Yun Lim, Young Suk Choi, Hye Rim Lee, Hye Min An). **Authors’ contributions** Ji Yun Lim is the first author, performed the experiments, and was responsible for the overall writing and revision of the manuscript. Young Suk Choi, Hye Rim Lee, and Hye Min An are co-authors, analyzed the result data and performed parts of the experiments. Young Koo Lee is a corresponding author, played a role in interpreting the results, and gave final approval.

## Abstract

Red ginseng is an immune-enhancing compound that exhibits anti-inflammatory action. The ginsenoside Rg1, an ingredient of red ginseng, has been shown to play an important role in tumor suppression, wound healing, and angiogenesis. This study evaluated the effects of red ginseng extract and Rg1 in a diabetic wound model. Diabetes was induced with streptozotocin (STZ) in 8-week-old male Institute of Cancer Research (ICR) mice weighing 30–35 g. A full-thickness skin defect was treated by applying a dressing every 3 days. The mice were divided into three groups. Group 1 was administered an extract of red ginseng (10 mg/kg/d, *n* = 27, oral) and group 2 was administered Rg1 (10 mg/kg/d, *n* = 27, oral). Group 3 was a control group treated with phosphate-buffered saline (0.3 mL/kg/d, *n* = 27, oral). Red ginseng extract and Rg1 were orally administered to mice daily for 10 days following injury in groups 1 and 2, respectively. Both increased mRNA and protein levels of vascular endothelial growth factor (VEGF) and transforming growth factor (TGF)-β1 compared to controls. In addition, the wounds of animals in the Rg1 group were significantly smaller between days 7 and 10 (*p* < 0.05). VEGF and TGF-β1 were not expressed in diabetic mice in the control group. Both red ginseng extract and Rg1 promoted the production of VEGF and TGF-β1, which are important in wound healing. Our results for Rg1 suggest its potential to promote diabetic wound healing by stimulating the production or activity of VEGF and TGF-β1 factors involved in the wound healing process.

## Introduction

By the year 2035, it is expected that 592 million people will suffer from diabetes, making it one of the most important medical issues globally [1]. Diabetes is divided into two types: type 1 diabetes (T1D) is characterized by impaired pancreatic β-cell function resulting in insulin deficiency and chronic hyperglycemia while type 2 diabetes (T2D) is characterized by insulin resistance and hyperglycemia [2]. Chronic hyperglycemia causes deterioration of blood vessels and nerves, resulting in cardiovascular diseases and neuropathy [3, 4]. Additional complications, such as diabetic wounds, impair the four stages of wound healing (e.g., hemostasis, inflammation, proliferation and remodeling), delaying the healing process [5]. As a result, chronic wounds in diabetic patients damage wounds and surrounding tissues due to excessive inflammatory reactions, and delay wound healing through delayed expression of growth factors. In serious cases, this results in loss of the affected limb or death [6, 7]. However, a detailed understanding of the delayed healing of diabetic wounds is lacking and current forms of treatment such as oral antidiabetics and insulin injection, have many unwanted side effects, such as hypertension, weight gain, and endothelial cell damage [8, 9]. This has stimulated the search for safe and effective treatments. *Panax ginseng* is one of the oldest traditional herbs and it contains approximately 32 species of ginsenosides [10]. In recent years, ginsenosides have been demonstrated to exhibit many properties, including immune-stimulating activity and anticancer, anti-inflammatory, antiallergic, antihypertensive, and antidiabetic effects [11]. Ginsenosides with antidiabetic effects include Rg1, Rg3, Rg5, Rb1, Rb2, and Rb3, and ginsenosides have been used as an adjuvant for the treatment of diabetes because they exhibit antidiabetic effects [12]. Of the many types of saponins, the ginsenoside Rg1 reduces intestinal glucose uptake by inhibiting the expression of the Na^+^/glucose cotransporter 1 (SGLT1) gene responsible for glucose uptake in intestinal epithelial cells [13]. Rg1 has been proposed to reduce oxidative stress and cardiomyocyte death and prevent cardiovascular damage in diabetic mice [14]. Rg1 is also excellent for skin regeneration [15]. Representative factors involved in wound healing include transforming growth factor (TGF)-β, fibroblast growth factor (FGF), and vascular endothelial growth factor (VEGF). VEGF plays an important role in wound healing, and participates in endothelial migration, proliferation, and granulation tissue and blood vessel formation. TGF-β promotes wound healing, fibroblast proliferation, and expression of major components of the extracellular matrix such as fibronectin, collagen I and III, and VEGF [16]. Although wounds heal as the expression of these growth factors gradually increases, the roles of these factors throughout the healing process have not been fully clarified [17]. The use of Rg1 in wound healing and angiogenesis is being actively studied [18,19], although neither its effect nor that of red ginseng on diabetic wounds is known. Therefore, the purpose of this pilot study was to evaluate the effects of red ginseng extract and Rg1 in a streptozotocin (STZ)-induced type 1 diabetic mouse wound model. The main hypothesis of this pilot study was that the expression of VEGF and TGF-β1 in animal models treated with red ginseng extract and Rg1 is important in wound healing.

## Materials and methods

### Animal model

Male Institute of Cancer Research (ICR) mice aged 6 to 8 weeks (30–35 g) were purchased from Orient BIO Inc. (Seongnam, Korea). All mice were allowed to adjust to the environment for 1 week prior to the experiment. The mice were housed in cages with a 12 h/12 h light/dark cycle at 23 ± 2°C and 50 ± 20% humidity with freely available water and rodent chow (LabDiet 5L79^®^; Orient BIO Inc.). The experimental design was approved by the Institutional Animal Care and Use Committee of Soonchunhyang University Medical School (SCHBC-Animal-2016–03).

### Diabetic wound model using STZ

Diabetes was induced by intraperitoneal (IP) injection of 100 mg/kg STZ (S0130; Sigma-Aldrich, St. Louis, MO, USA) in 0.1 M citrate buffer (pH 4.5) to 81 mice following a 6 h fast. After 24 h, blood was collected from the tail vein and blood glucose levels were measured using the Accu-Chek^®^ system (Roche Diagnostics, Mannheim, Germany). Diabetes was defined as a blood glucose level greater than 200 mg/dL after STZ injection. The mice were anesthetized by IP injection of zolazepam/tiletamine (Zoletil 50^®^; Virbac, Carros, France) and xylazine (Rompun^®^; Bayer Korea, Seoul, Korea). The hairs on the back of the mice were shaved, the exposed skin was cleansed with 70% ethanol, and a full-thickness skin defect wound of 8 mm in diameter was made using a sterile skin biopsy punch (BP-80F; Kai Industries, Gifu, Japan). The methicillin-resistant *Staphylococcus aureus* (MRSA) standard strain (ATCC^®^; 43300 MINIPACK™, Manassas, VA, USA) was incubated overnight at 37°C on Mueller – Hinton agar plates. MRSA was diluted in sterilized saline to adjust to McFarland 0.5 standard turbidity and applied to the skin defects. Vaseline (Samhyun Pharmaceutical, Seoul, Korea) was applied to all wounds using a gauze pad to prevent the wound from drying. Then the wounds were covered with Opsite-film (Opsite Flexifix^®^; Smith & Nephew Medical Ltd., Hull, UK) dressings. The dressings were replaced every 3 days.

### Experimental design for red ginseng extract and Rg1 treatment

STZ-induced diabetic mice were orally administered 10 mg/kg of red ginseng extract or Rg1 once daily for 10 days after induction of injury. Concentration and date were selected according to Yu *et al.* [20]. Red ginseng extract (Korean Red Ginseng Extract Gold^®^; The Korean Ginseng Research Institute, Daejeon, Korea) and the ginsenoside Rg1 (purity > 98%; 22427– 39–0; Abcam, Cambridge, UK) were dissolved in phosphate-buffered saline (PBS). The STZ-induced diabetic mice (*n* = 81) were randomly divided into three different treatment groups as follows: PBS (0.3 mL/kg/d, *n* = 27, oral), red ginseng extract (10 mg/kg/d, *n* = 27, oral), and Rg1 (10 mg/kg/d, *n* = 27, oral). Treatments were administered orally once daily for 10 days following induction of injury (Figure 1).

**Figure 1.**
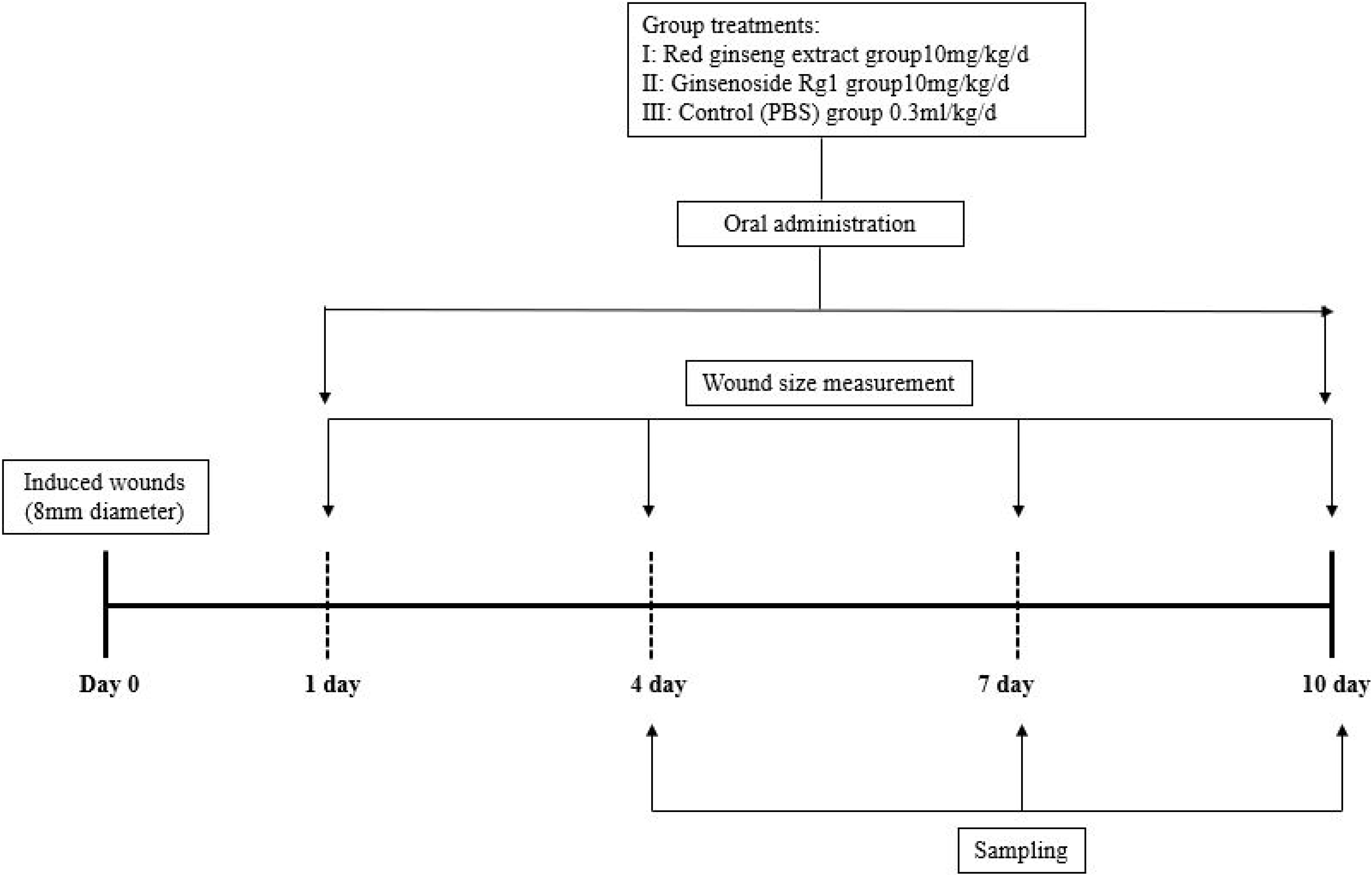
Protocol for wound induction. STZ-induced diabetic mice were orally administered red ginseng extract (10 mg/kg) and Rg1 (10 mg/kg). Wounds were made on day 0 and measured on days 1, 4, and 7. Mice were sacrificed on days 4, 7, and 10. Mice were randomly divided into three groups according to oral-administration treatment. G1: red ginseng extract treatment group (10 mg/kg/day, *n* = 27); G2: Rg1 treatment group (10 mg/kg/day, *n* = 27); G3: control (PBS) group (0.3 mL/kg/day, *n* = 27). STZ: streptozotocin, Rg1: ginsenoside Rg1.

### Measurement of wound area

Images of the skin wound were obtained on days 1, 4, 7, and 10 and the wound size was measured using the UTHSCSA image tool (version 3.0; Microsoft Corporation, Redmond, WA, USA). The wound size on the first day after wound induction was defined as 100% and wound sizes measured on days 4, 7, and 10 are expressed as a percentage of wound size at day 1.

### Histological analyses of diabetic wound tissue

Wound tissues were compared in the three groups by euthanizing the mice at days 1, 4, 7, and 10 after injury. The skin tissues were harvested to include the entire wound, fixed for 12 h in 4% paraformaldehyde prepared in PBS (pH 7.4; Santa Cruz Biotechnology, Dallas, TX, USA), washed with PBS at room temperature for 6 h, and processed overnight in an automatic tissue processor. Sections of 4 μm thickness were cut, placed on coated glass slides (5116–20F; Muto Pure Chemicals Co., Ltd., Tokyo, Japan), and deparaffinized before staining with hematoxylin and eosin (H&E; Gills hematoxylin, 1% eosin; Muto Pure Chemicals Co., Ltd.). Three pathologists blinded to the three groups scored the tissues according to Tables 1 and 2 based on light microscopy (DXM1200; Nikon, Tokyo, Japan) images of the sections at 200× magnification. These images were assessed for granulation tissue area, inflammatory cells, connective tissue, and damage to the dermal layer.

**Table 1.**
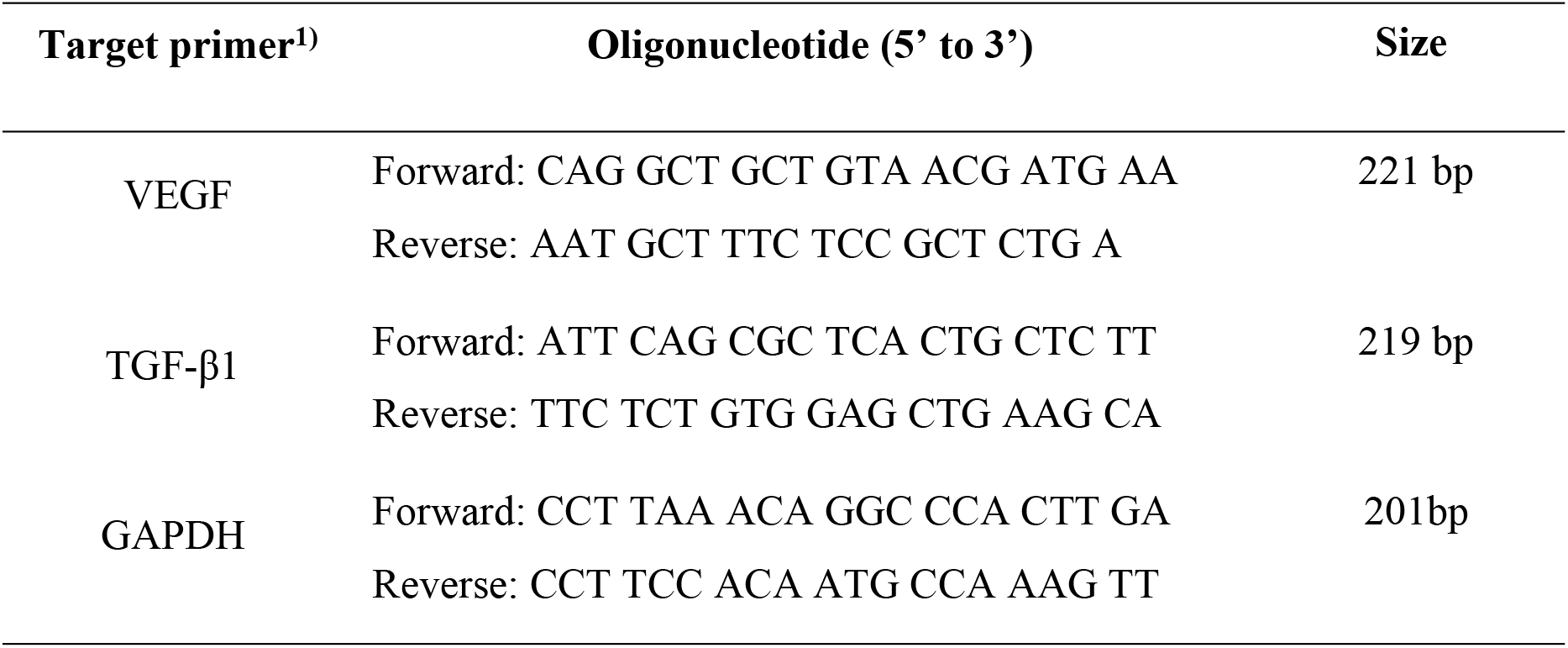
Reverse transcription-polymerase chain reaction primer sequence. VEGF, vascular endothelial growth factor; TGF-β1, transforming growth factor-β1; GAPDH, glyceraldehyde-3-phosphate dehydrogenase. ^1)^ All primers were designed directly and purchased from Macrogen Corporation (Seoul, Republic of Korea). The primers for VEGF, TGF-β1, and GAPDH used in this experiment are shown in the table and reverse transcription– polymerase chain reaction was performed using each primer.

**Table 2.** Degree of granulation tissue of formation score. The degree of granulation tissue formation was scored in mouse skin tissue. Hematoxylin and eosin staining were used to evaluate the degree of granulation formation histologically.

| Judgment element | Standard | Score |
| --- | --- | --- |
| Granulation tissue of formation | Not formation | 0 |
|  | More than 2/3 of defect area | 1 |
|  | 1/3 to 2/3 of defect area | 2 |
|  | Less than 1/3 of defective area | 3 |

### Expression of VEGF and TGF-β1 in diabetic wound tissue by immunohistochemistry

The paraffin-embedded wound tissue blocks were sectioned, placed on slides, and incubated at 60–70°C for 30 min. The sections were deparaffinized in a graded series of xylene and ethanol (100%, 95%, 80%, and 70%) and treated with an endogenous peroxidase blocking agent, which then was diluted with methanol (99%) and a 1.4% hydrogen peroxide solution (FW = 34.01). After 30 min, the slides were washed with 1× Tris-buffered saline (TBS) and sections were blocked with 1.5% normal serum (PK-6102, horse serum, mouse IgG or PK-6101, goat serum rabbit IgG; Vector Laboratories, Burlingame, CA, USA) for 1 h followed by incubation overnight at 4°C with anti-VEGF monoclonal antibody (SC-7269, 1:500 dilution; Santa Cruz Biotechnology) and anti-TGF-β1 polyclonal antibody (ab92486, 1:1000 dilution; Abcam) in blocking buffer. Then the samples were washed with 1× TBS followed by incubation for 1 h at room temperature with a biotinylated secondary antibody (PK-6102, horse serum, mouse IgG or PK-6101, goat serum rabbit IgG; 1:3000 dilution; Vector Laboratories) in blocking buffer, and washed again with 1× TBS. Incubation with the VECTASTAIN Elite avidin-biotin complex (PK-6102 or PK-6101; Vectastain®; Vector Laboratories) was used to increase the target antigen-antibody reaction during a 30 min incubation. Then the sections were stained with the chromogen diaminobenzidine (C02-100; Liquid DAB + Chromogen Kit; Golden Bridge International Inc., Mukilteo, WA, USA) for 5–6 min followed by hematoxylin for 1 min before they were covered with a coverslip and evaluated using a digital microscope (DXM1200; Nikon) at 200× magnification.

### Reverse transcription–polymerase chain reaction (RT-PCR)

Total RNA was isolated from wounded skin tissue samples using TRIzol Reagent^®^ (TR118; Molecular Research Center, Cincinnati, OH, USA). mRNA was isolated using chloroform and isopropanol (C2432, I9516; Sigma-Aldrich). After precipitation with isopropanol, the RNA pellet was dissolved in distilled water treated with diethyl pyrocarbonate (DEPC; WR2004; Biosesang, Sungnam, Korea). Total RNA was quantified to 3 μg to react 1 μg/μL random hexamer (C1181; Promega, Madison, WI, USA) at 70°C for 5 min, then incubated at 4°C for 5 min. Each reaction contained 1 μL of 10 mM dNTPs, 1 μmol/L MgCl_2_, 1 μg/μL RNaseOUT, 1 μg/μL 5×reaction buffer, 1 μL GoScript reverse transcriptase (A5003; Promega), and 7 μL RNase-free double-distilled H2O (N2511; Promega) and was incubated at 25°C for 5 min, followed by incubation at 42°C for 1 h and 70°C for 15 min. The VEGF, TGF-β1, and glyceraldehyde-3-phosphate dehydrogenase (GAPDH) primer gene sequences were as follows: VEGF forward primer, 5’- CAG GCT GCT GTA ACG ATG AA -3’; VEGF reverse primer, 5’- AAT GCT TTC TCC GCT CTG A -3’; TGF-β1 forward primer, 5’- ATT CAG CGC TCA CTG CTC TT-3’; TGF-β1 reverse primer, 5’- TTC TCT GTG GAG CTG AAG CA-3’; and GAPDH forward primer, 5’- CCT TAA ACA GGC CCA CTT GA-3’; GAPDH reverse primer, 5’ - CCT TCC ACA ATG CCA AAG TT-3’ (Table 1). All primers were ordered from Macrogen Oligo (Seoul, Korea). PCR was performed using 1 μL cDNA in a mixture with 2× PCR premix (LGT-1212; Lugen Science Co., Seoul, Korea). The conditions for VEGF included initial denaturation at 95°C for 5 min, melting at 95°C for 30 s, annealing at 60°C for 1 min, and elongation at 72°C for 30 s. For TGF-β1 and GAPDH, annealing was performed at 58°C for 30 s. PCR products were visualized using a bioimaging system (C280; Azure Biosystems, Dublin, CA, USA) after electrophoresis on a 1% agarose gel containing Safe Shine Green DNA Staining Solution (GC6051; Biosesang). GAPDH was selected for normalization of specific primer target bands, and density measurements of the target bands were analyzed with ImageJ software (National Institutes of Health, Bethesda, MD, USA).

### Western blot analysis

The skin tissue was homogenized using complete EDTA free protease inhibitor cocktail (11836170001; Roche Diagnostics GmbH), phosSTOP phosphatase inhibitor (30498800; Roche Diagnostics GmbH), and RIPA buffer (R4200; GenDEPOT, Barker, TX, USA). Dissolved protein was analyzed for protein concentration by micro-scale BCA assay (Micro BCA Protein Assay Kit #23225; Thermo Fisher Scientific Inc., Waltham, MA, USA). Lysates measured at 50 μg were separated using 10% sodium dodecyl sulfate-polyacrylamide electrophoresis (SDS-PAGE) and transferred to a polyvinylidene difluoride membrane (10600023; GE Healthcare Life Sciences, Chalfont, UK). After blocking with 5% skim milk (232100; Becton Dickinson, Franklin Lakes, NJ, USA) at 4°C for 24 h, the membranes were incubated with anti-VEGF monoclonal antibody (SC-7269; 1:500 dilution; Santa Cruz Biotechnology), anti-TGF-β1 polyclonal antibody (ab92486; 1:1000 dilution; Abcam), and β-actin rabbit antibody (4970S; 1:5000 dilution; Cell Signaling, Beverly, MA, USA) at room temperature for 2 h followed by the appropriate secondary antibody (SC-2005; 1:3000 dilution; goat anti-mouse or SC-2030; 1:3000 dilution; goat anti-rabbit; Santa Cruz Biotechnology) for 1 h. Bands of the target size were visualized by enhanced chemiluminescence (ECL 2232; GE Healthcare Life Sciences) and a bioimaging system (C280; Azure Biosystems). The band intensities were normalized with β-actin and the density measurements were analyzed with ImageJ software (National Institutes of Health).

### Statistical analysis

The data were analyzed using SPSS version 22.0 (SPSS Inc., Chicago, IL, USA) and the Mann– Whitney U test. All *p*-values less than 0.05 were considered statistically significant.

## Results

### Wound size measurement

In each group, the wound size decreased over time (Figure 2A). On day 4, there was larger decrease in wound size in the two treatment groups compared to the control group. From days 7 to 10, wound size was significantly decreased in the Rg1 group (*p* < 0.05). These results showed that wound contraction was the highest and wound healing was the fastest in the Rg1 group (Figure 2B).

**Figure 2.**
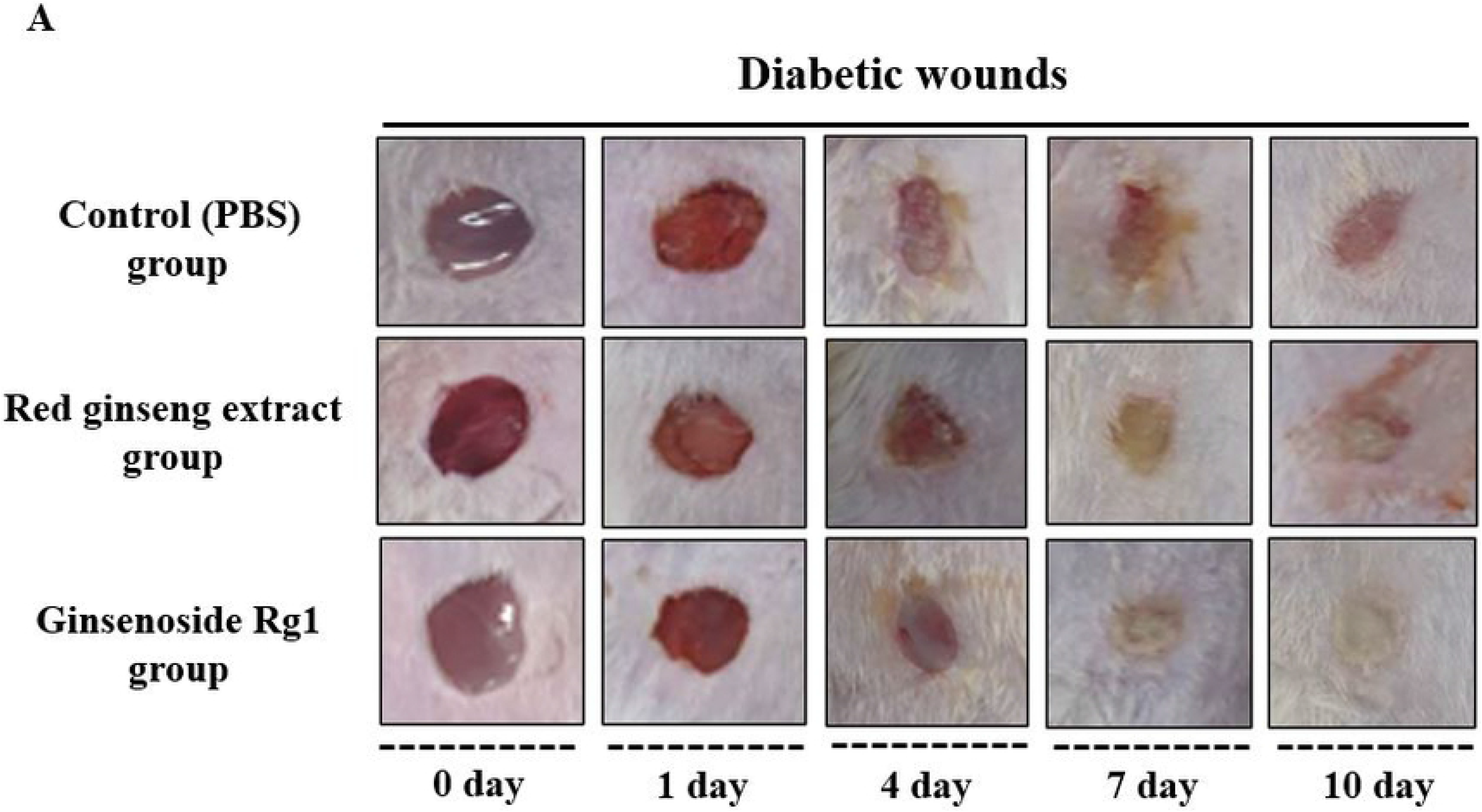
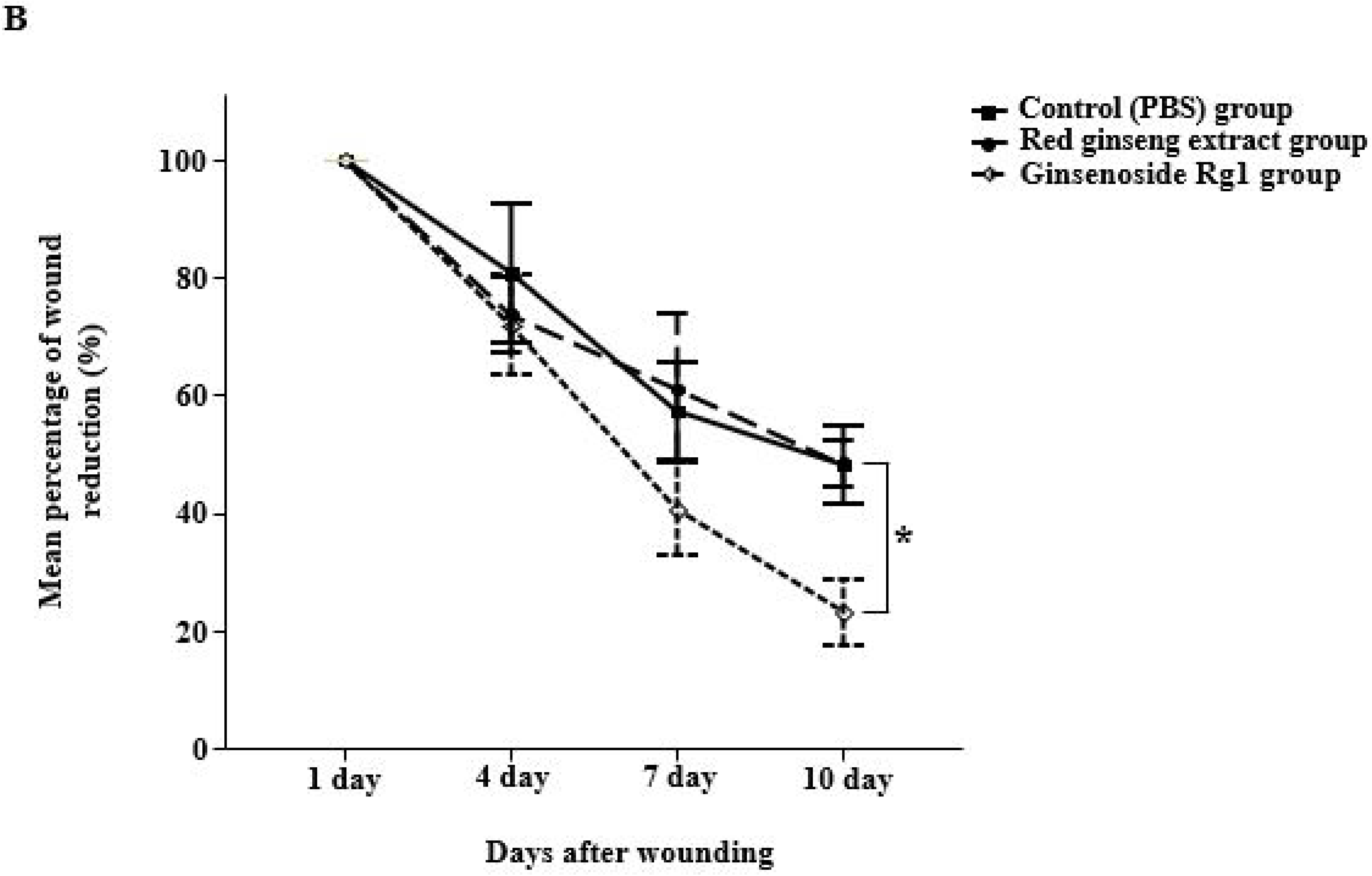
Changes in wound size by date in MRSA-infected diabetic rats. (A) Images of wound size on different dates. (B) Wound size measurements for each group. At each point, the wound healing rate changed compared to the first day of injury. Data are expressed as a percentage by measuring the wounds of the mice in each group. Data are expressed as mean ± standard deviation (*n* = 9). MRSA: methicillin-resistant *Staphylococcus aureus.*

### Histology and immunohistochemistry analyses

The central portion of the wounded tissues was collected to evaluate granulation tissue formation, inflammatory cell infiltration, and growth factor expression. The wound healing effects of the red ginseng extract and Rg1 groups were evaluated histologically. Samples of all groups exhibited histological patterns of dermal tissue (consisting of three layers of epidermis, dermis, and subcutaneous tissue). H&E and immunohistochemistry (IHC) staining were performed (Figure 3A–C). The degree of granulation and inflammatory cell infiltration was not different in the three groups at day 4. However, at day 10, granulation tissue was formed in the Rg1 group whereas the control group remained unchanged (*p* < 0.05). On the other hand, inflammatory cell infiltration in the Rg1 group was significantly lower at day 10 compared to the other two groups (*p* < 0.05) (Figure 3D, E). Expression of VEGF and TGF-β1 was confirmed by IHC staining; VEGF was mainly present in granulation tissues and TGF-β1 was observed on the wound surface.

**Figure 3.**
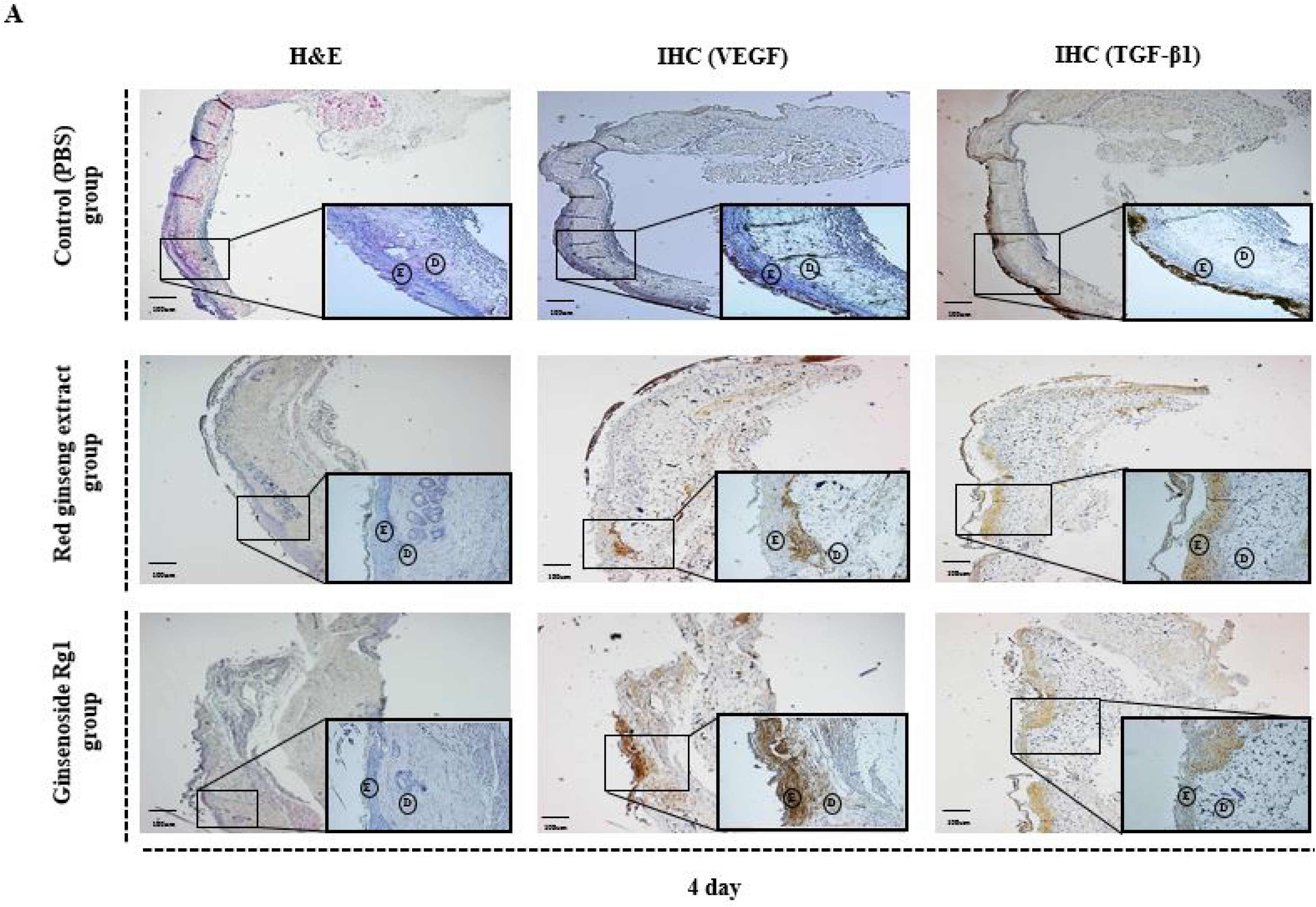
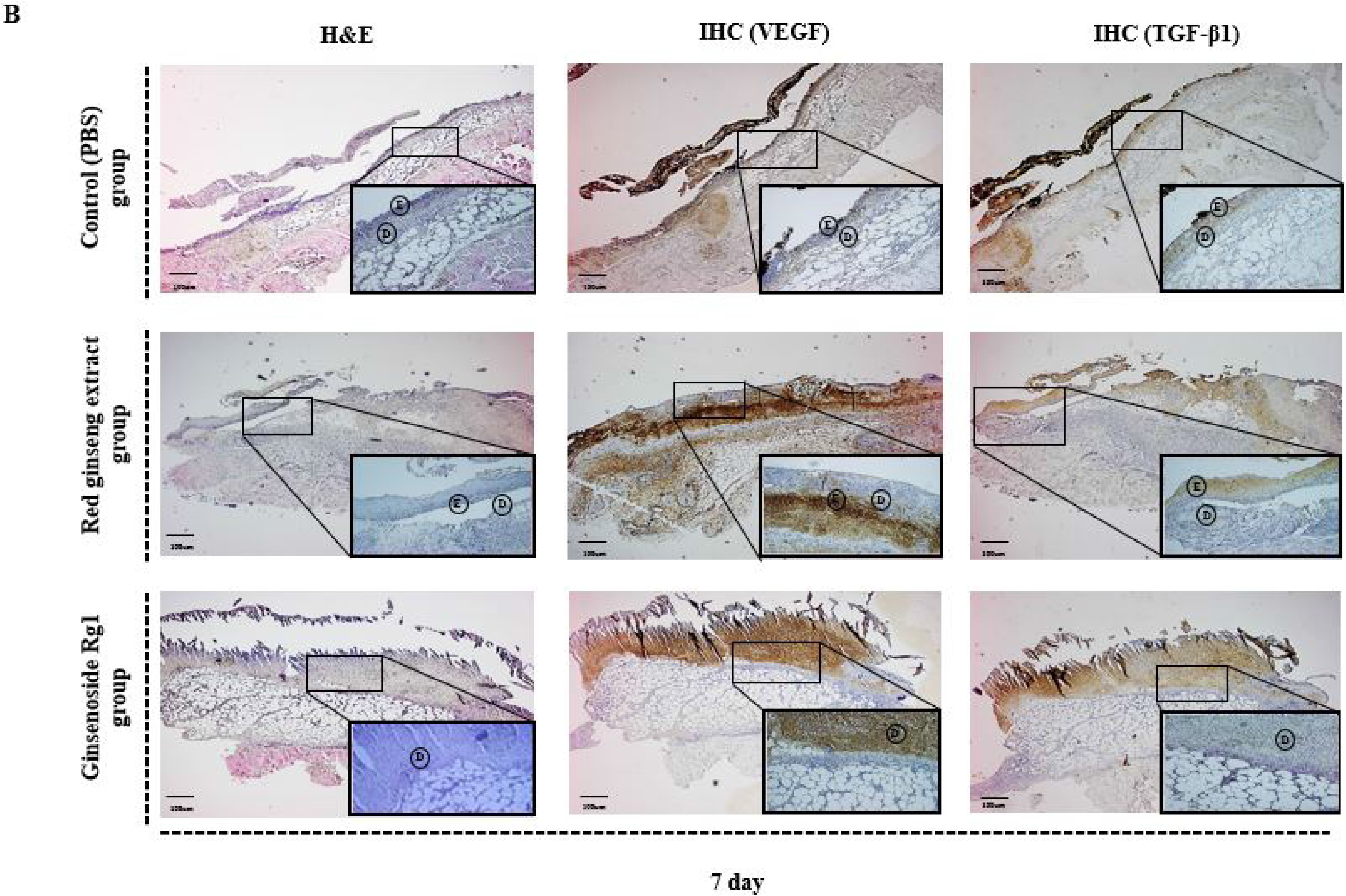
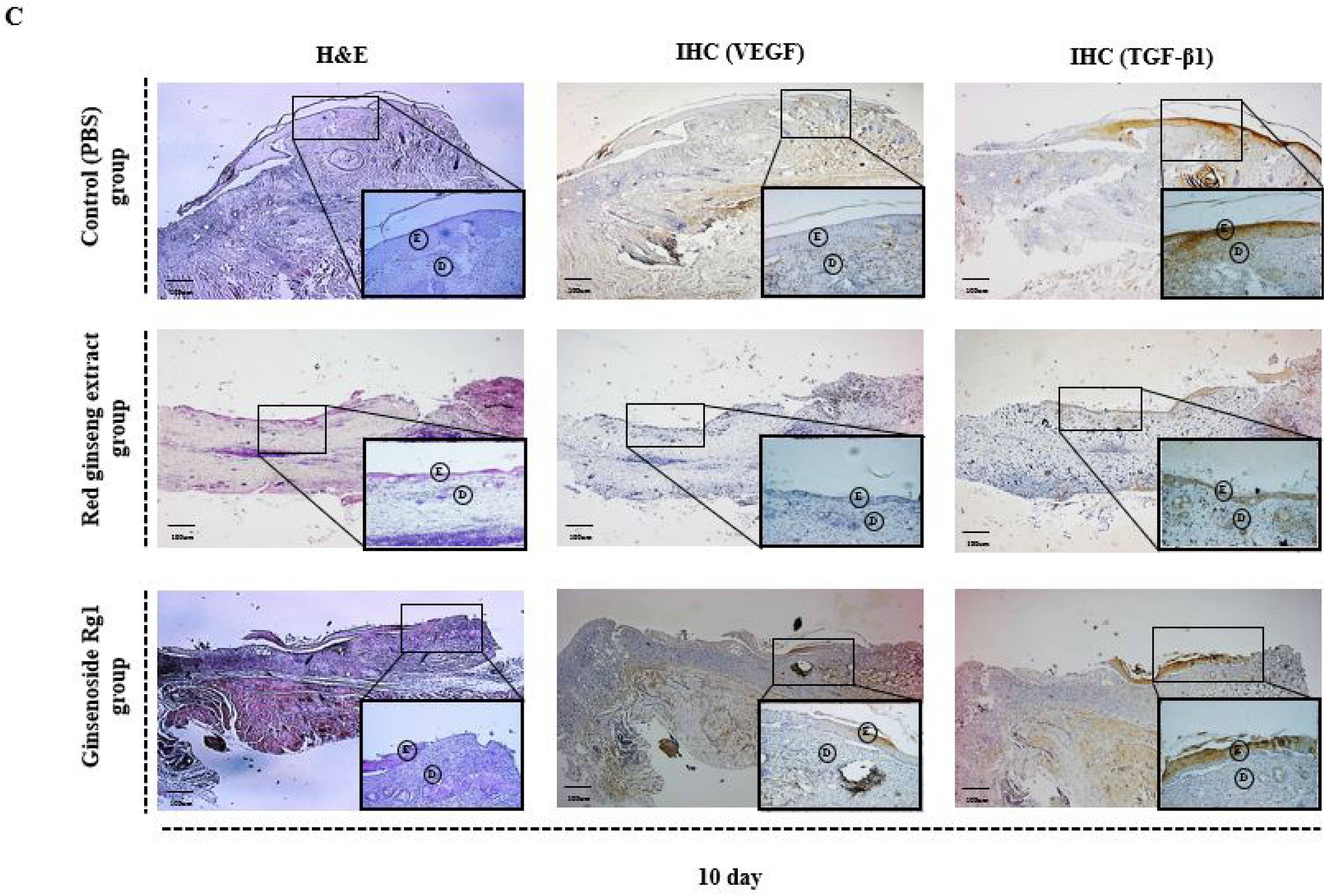
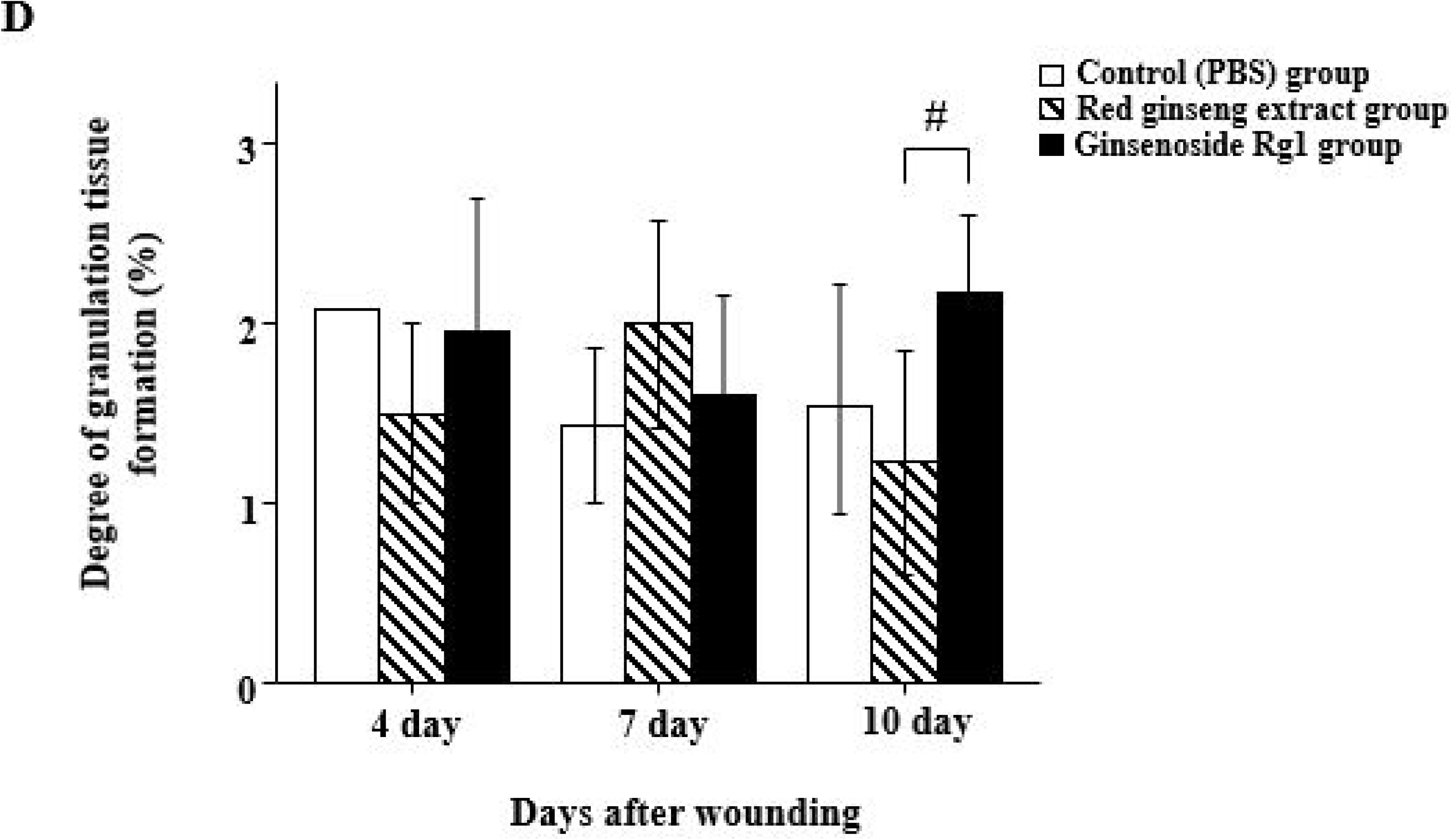
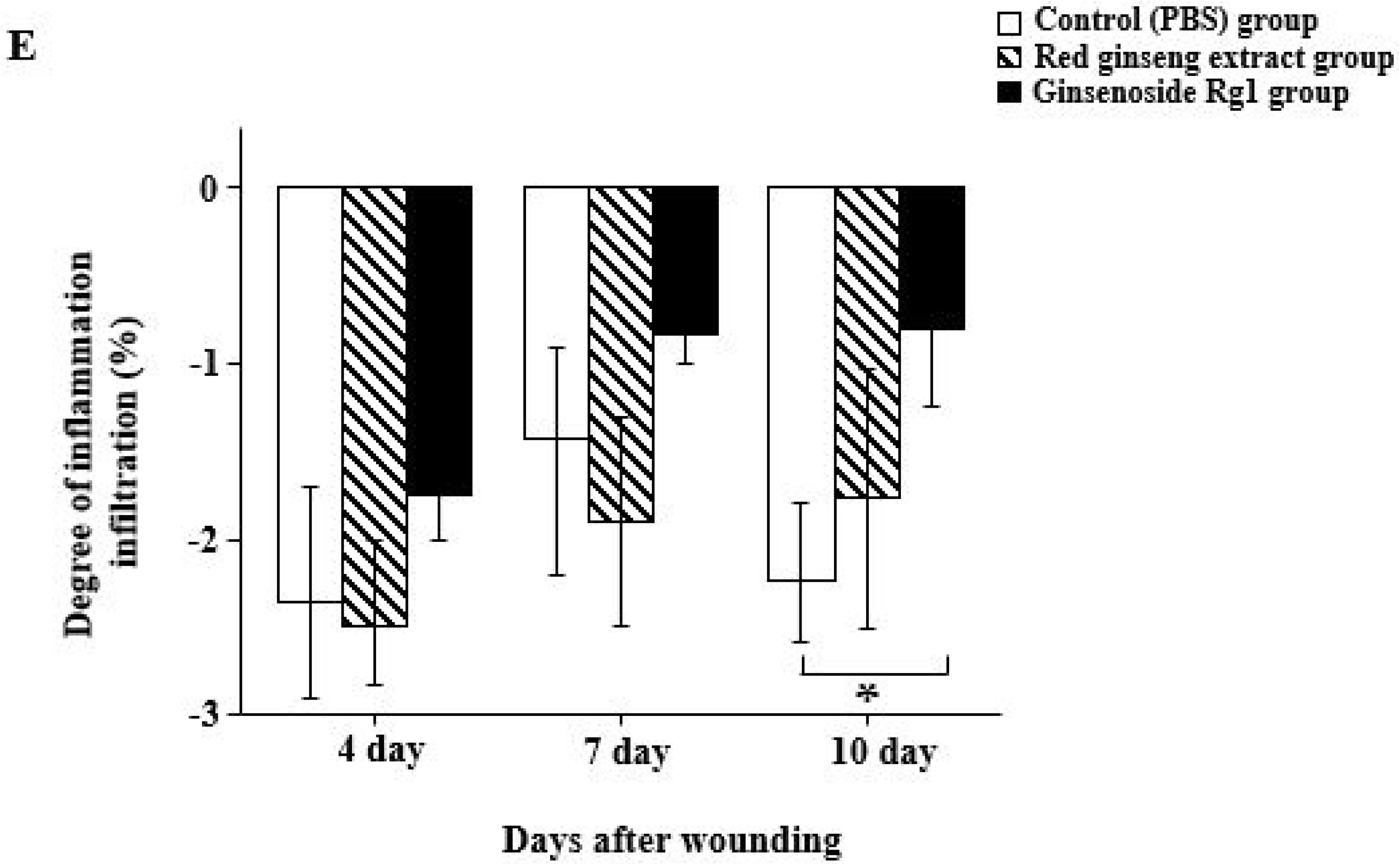
Histology and IHC analysis. For histological analysis, the slides corresponding to days 4, 7, and 10 for each group were stained with hematoxylin and eosin (H&E). IHC staining of VEGF and TGF-β1 (important factors for wound healing) was observed. (A) Tissues from mice of each group sacrificed 4 days after injury (H&E stain in the left panels, VEGF antibody IHC stain in the middle panels, and TGF-β1 antibody IHC stain in the right panels). (B) Tissues from mice of each group sacrificed 7 days after injury. (C) Tissues from mice of each group sacrificed 10 days after injury. The original magnifications of 200× and 400× (bottom panel) are displayed. (^*^) Position of the granulation tissue and wound. (D) The degree of granulation tissue formation and infiltration of inflammatory cells were scored to objectify as H&E stain (Tables 2 and 3). An objective scoring table shows that repeated scoring of three times was performed by blind test. IHC staining shows hematoxylin (blue) and VEGF and TGF-β1 antibody (brown). Mice were divided into three groups (red ginseng extract group, Rg1 group, and the PBS-treated group as a control). *p < 0.05 compared to the control group and # p < 0.05 compared to the treatment group. E: epidermis, D: dermis, TGF: transforming growth factor, VEGF: vascular endothelial growth factor.

**Table 3.** Degree of inflammatory cells infiltration score. The degree of infiltration of inflammatory cell was scored in mouse skin tissue. Hematoxylin and eosin staining was performed histologically to evaluate the degree of infiltration of inflammatory cell.

| Judgment element | Standard | Score |
| --- | --- | --- |
| Inflammatory cell of infiltration | No necrosis and inflammation | 0 |
|  | Not clusters and only exist in cell form | -1 |
|  | When small clusters are formed and inflammatory cells are visible | -2 |
|  | When large clusters are formed and many inflammatory cells are visible | -3 |

### Expression of VEGF and TGF-β1 by RT-PCR

We investigated the effects of wound healing on mRNA expression of VEGF and TGF-β1 in a diabetic wound model. GAPDH was used to normalize the expression level of specific target bands (Figure 4A). At day 7, mRNA expression of VEGF and TGF-β1 was significantly increased in the red ginseng extract group (*p* < 0.05) and decreased at day 10. The expression of VEGF in the Rg1 group significantly increased on days 4–10 (*p* < 0.05). Expression of TGF-β1 was observed from day 4 in the red ginseng extract group but gradually increased on days 7–10 in the Rg1 group. In the control group, the expression of VEGF and TGF-β1 was delayed (Figure 4B–C).

**Figure 4.**
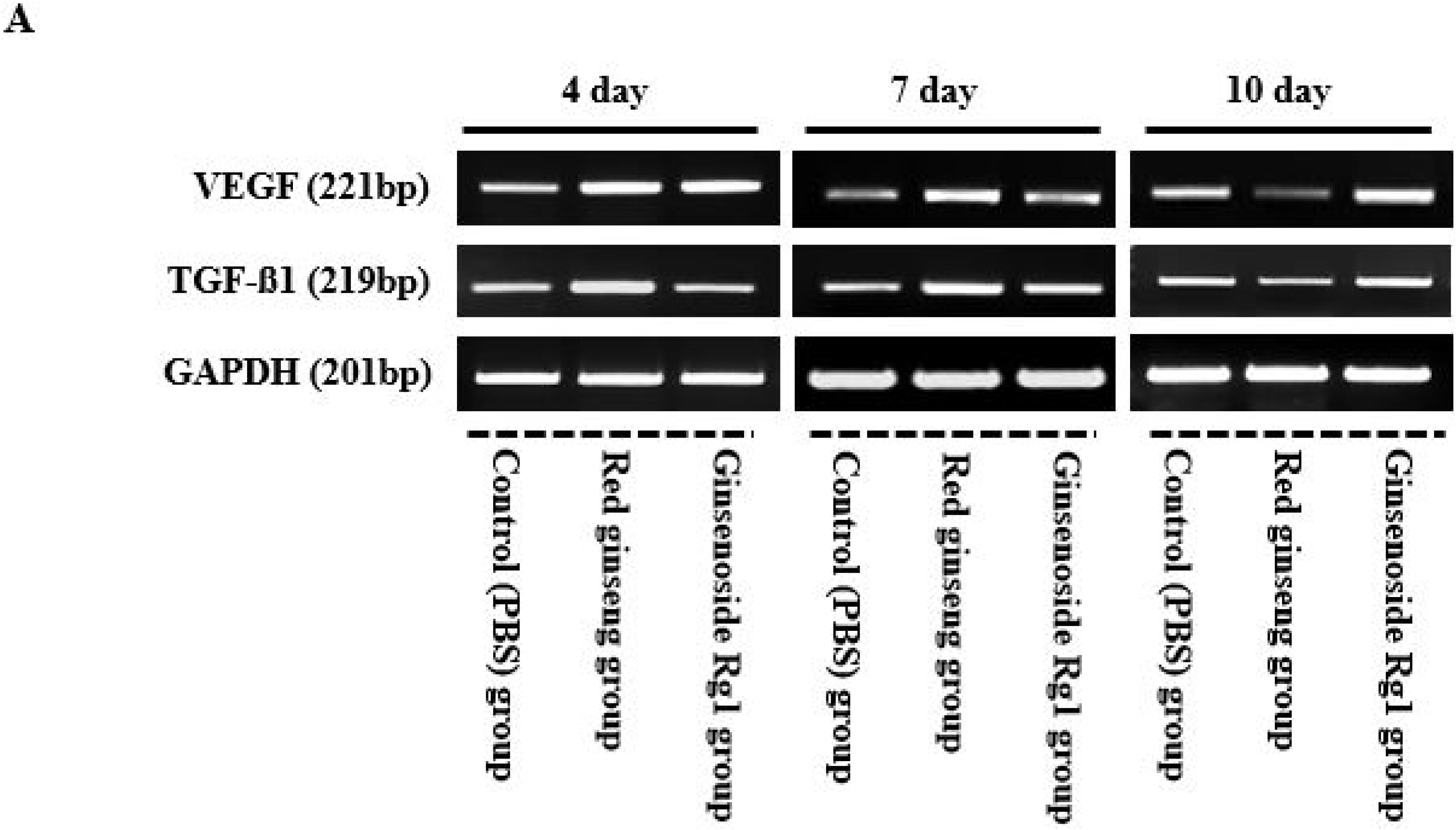
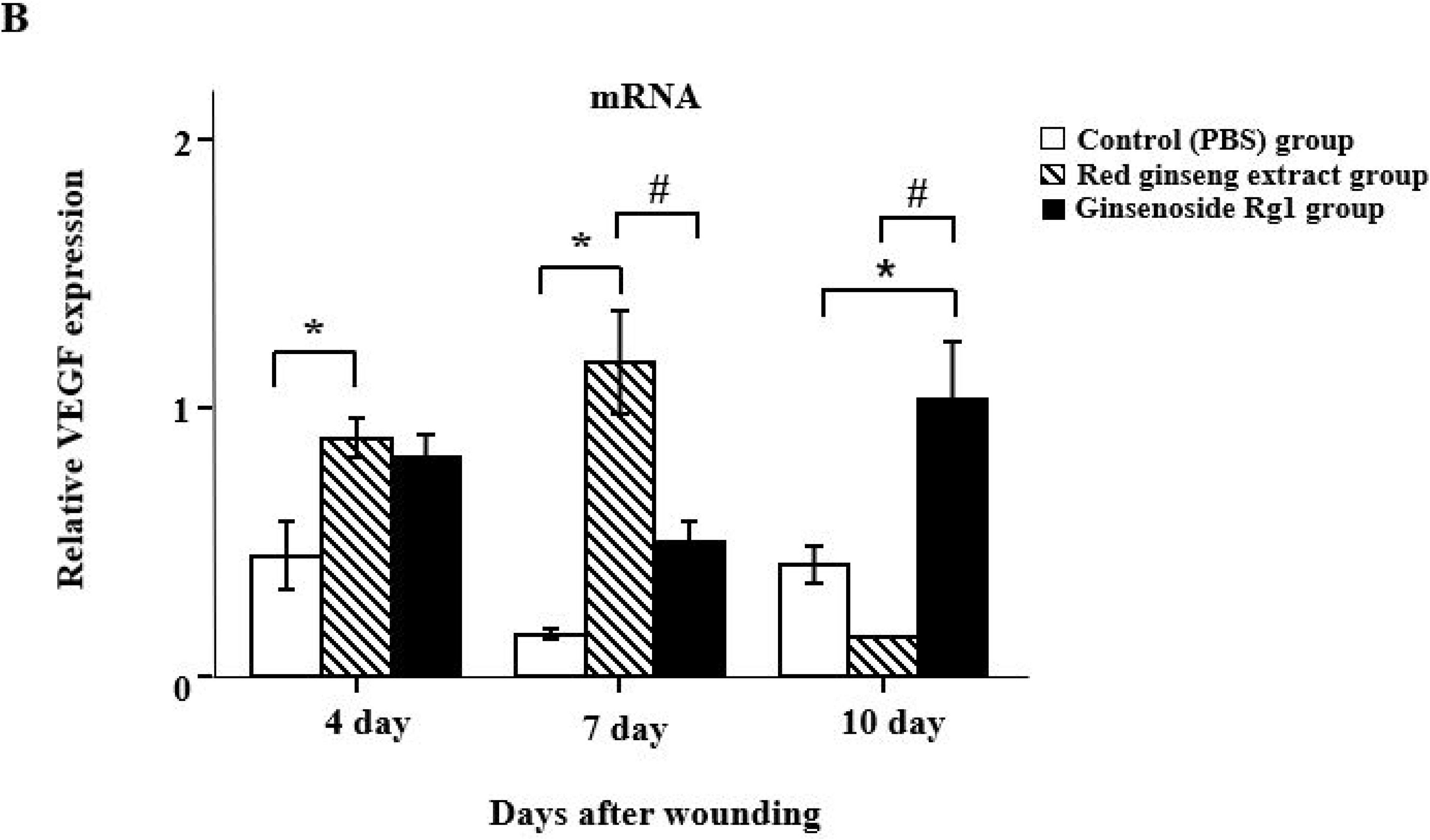
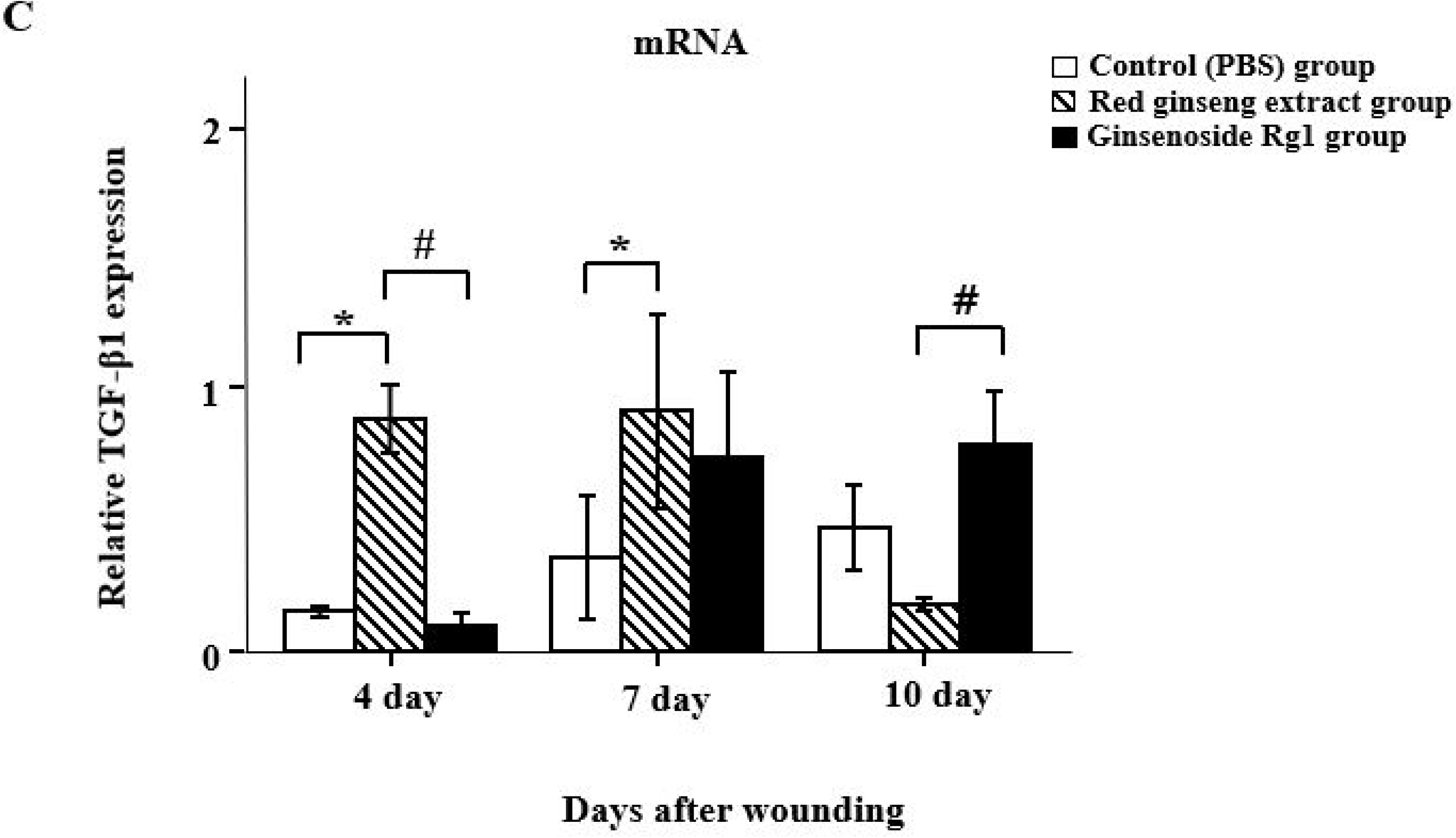
Effects of red ginseng extract and Rg1 on the expression of VEGF and TGF-β1 in diabetic wounds indicated by reverse transcription polymerase chain reaction (RT-PCR) analysis. (A) RT-PCR was performed for the tissue of diabetic mice using VEGF and TGF-β1 primers. GAPDH was used to normalize the intensity of the analyzed bands. (B–C) mRNA expression of VEGF and TGF-β1 increased significantly and then decreased during the wound-healing period (4–10 days after wound injury) in the red ginseng extract group; expression of VEGF and TGF-β1 increased gradually in the Rg1 group. Significance was determined using the Mann–Whitney U test. Mice were divided into three groups (red ginseng extract group, Rg1 group, PBS-treated group as a control group). *p < 0.05 compared to the control group and # p < 0.05 compared to the treatment group. TGF: transforming growth factor, VEGF: vascular endothelial growth factor, GAPDH: glyceraldehyde-3-phosphate dehydrogenase.

### Expression of VEGF and TGF-β1 by western blotting

Protein levels of VEGF and TGF-β1 were observed by western blotting; β-actin was used to normalize the expression level of specific target bands (Figure 5A). VEGF expression was significantly increased (*p* < 0.05) in the red ginseng extract and Rg1 groups at days 4–7, but decreased significantly at day 10 in the Rg1 group (*p* < 0.05). The expression of TGF-β1 was significantly higher on days 4–7 in the red ginseng extract group than in the other groups (*p* < 0.05) and decreased on days 7–10. In the control group, VEGF tended to increase over time, but TGF-β1 expression did not (Figure 5B–C).

**Figure 5.**
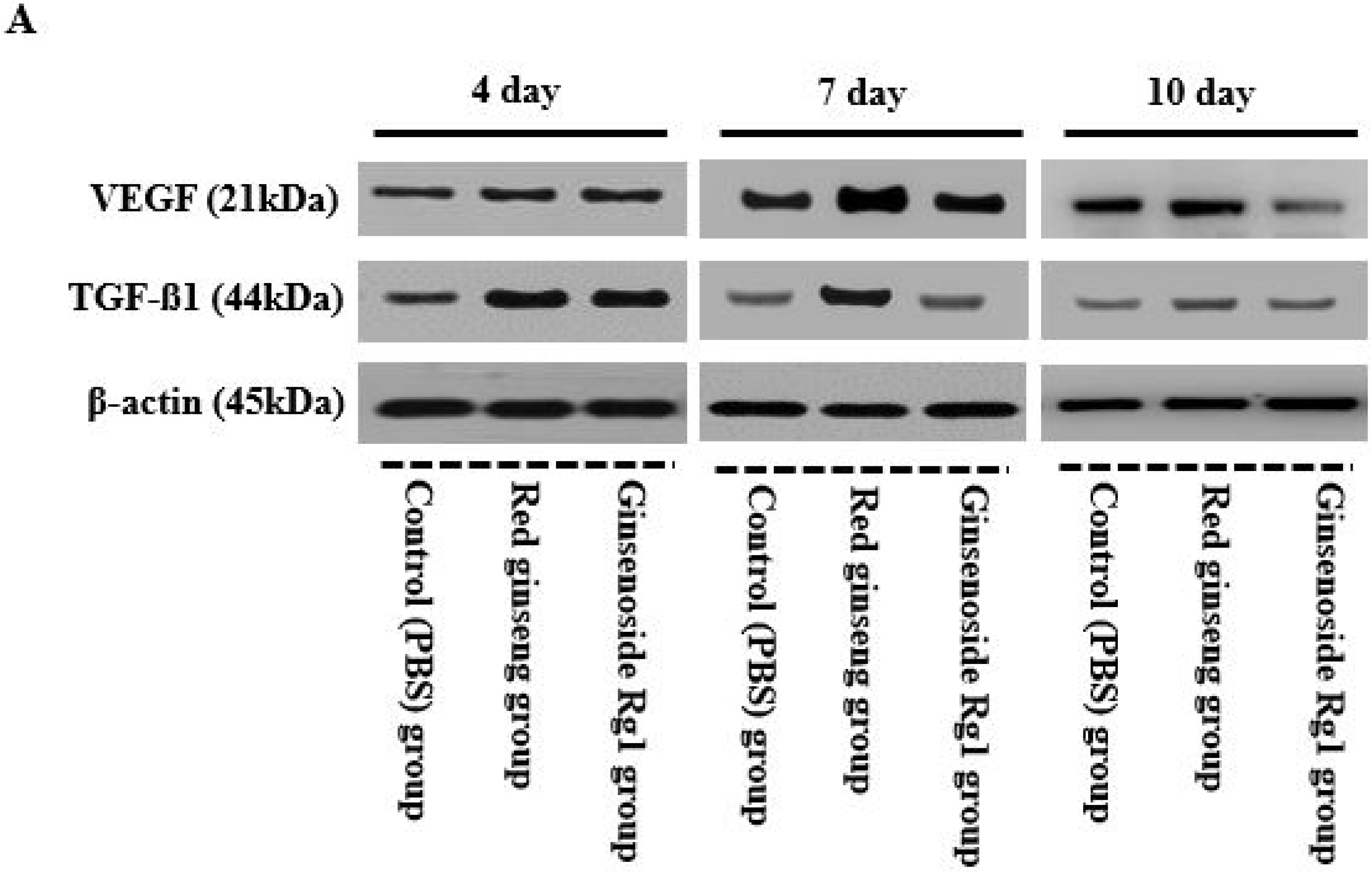
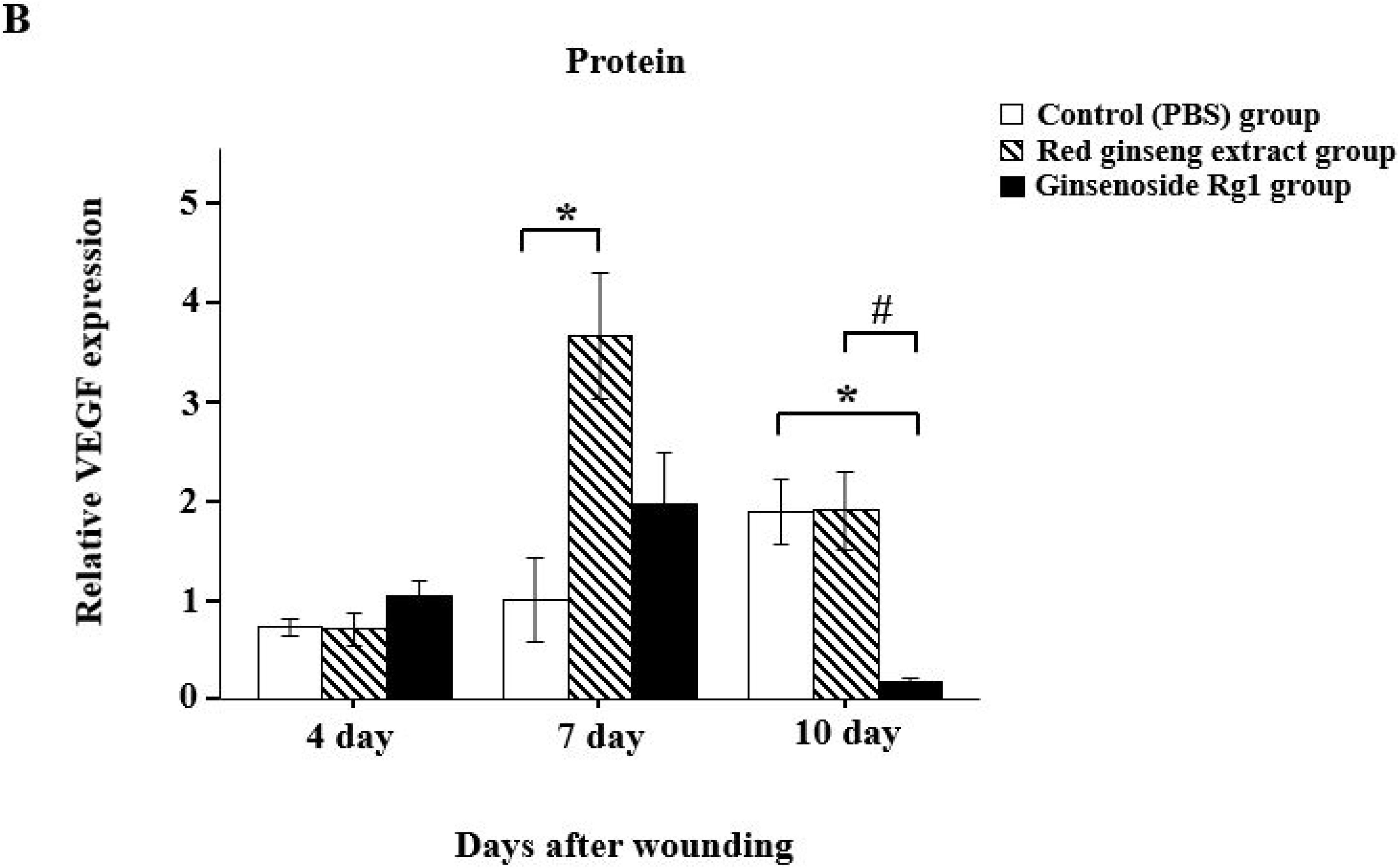
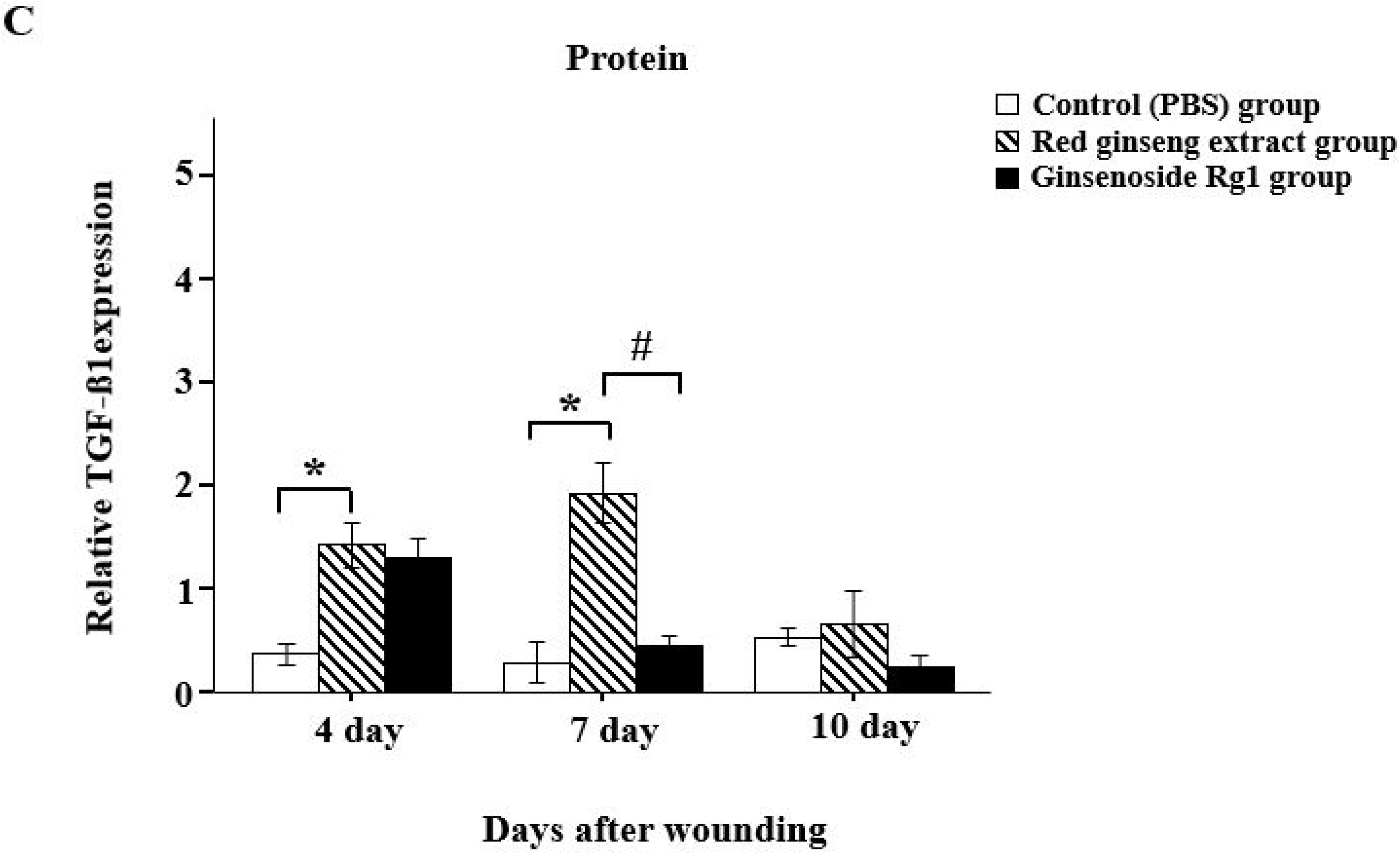
Effects of red ginseng extract and Rg1 on the expression of VEGF and TGF-β1 in diabetic wounds indicated by western blot analysis. (A) Immunoblotting was performed for diabetic mouse tissue using VEGF and TGF-β1 antibodies. β-actin was used to normalize the intensity of the analyzed bands. (B–C) Protein expression of VEGF and TGF-β1 increased significantly in the red ginseng extract and Rg1 groups during the wound healing period (4–10 days after injury) and then decreased with time. Significance was determined using the Mann– Whitney U test. Mice were divided into three groups (red ginseng extract group, Rg1 group, PBS-treated group as a control group). *p < 0.05 compared to the control group and # p < 0.05 compared to the treatment group. TGF: transforming growth factor, VEGF: vascular endothelial growth factor, β-actin: beta-actin.

## Discussion

This study indicates that administration of red ginseng extract and Rg1 can regulate the healing of diabetic wounds. Red ginseng extract and Rg1 have antidiabetic and wound healing effects on normal wounds [21, 22], but the effects on diabetic wound animal models have not been confirmed. This study showed that administration of red ginseng extract and Rg1 increases the expression of VEGF and TGF-β1, thereby promoting wound healing in an animal model of STZ-induced diabetic wounds (i.e., chronic wounds with delayed healing). Wound healing is generally divided into four stages: hemostasis, inflammation, proliferation, and remodeling. The first stage is hemostasis and clotting [23]. When this process begins, various immune cells such as platelets, neutrophils, and monocytes, as well as growth factors, such as platelet-derived growth factor (PDGF) and TGF-β, are expressed. Inflammatory processes produce neutrophils, macrophages, and inflammatory cytokines following tissue damage [24]. Some growth factors, such as PDGF, epidermal growth factor (EGF), and VEGF, are also expressed; however, in diabetic patients, due to this imbalance of cytokines, would healing does not follow the four general stages [25, 26], resulting in delayed healing. Rg1 not only lowers blood sugar but also increases the expression of factors involved in wound healing, thus promoting wound healing in mice [27, 28]. Kim *et al.* [29]. reported that RG promotes angiogenesis by stimulating VEGF and improves the proliferation of epidermal cells by upregulating cytokine expression in keratinocytes. These results suggest that RG accelerates the wound-healing process by increasing TGF-β and VEGF expression in the early stages of wound healing [30]. Diabetes is characterized by the overexpression of factors such as nuclear factor-κB as well as pro-inflammatory mediators, cytokines, and nitric oxide (NO), which increase intracellular oxidative stress due to insulin resistance and hyperglycemia, resulting in abnormal cells and inhibition of angiogenesis [31]. In addition, high glucose-derived reactive oxygen species affect the onset of diabetic complications [32]. Recent studies have shown that growth-factor regulatory defects in wounds in both diabetic models and diabetic patients exacerbate wound-healing disorders [33, 34]. Although prophylactic treatment with red ginseng contributes to wound healing by controlling the expression of VEGF and TGF-β1, excessive expression can lead to chronic inflammation and negatively affect wound healing [35, 36, 37]. Therefore, it is important to evaluate the inflammatory response in diabetic wounds and the underlying molecular mechanisms of growth factors. In the present study, we examined whether the administration of red ginseng extract and Rg1 affects expression of VEGF and TGF-β1 in a diabetic wound model. Our results provide evidence of the wound healing effects of red ginseng extract and Rg1 on the diabetic wound model. To determine whether red ginseng extract and Rg1 are effective for wound healing at mRNA and protein levels, we confirmed the expression of VEGF and TGF-β1 by RT-PCR and western blotting. Treatment with Rg1 in a diabetic wound model with delayed wound healing resulted in upregulated mRNA expression level of VEGF and TGF-β1 from the wound-healing inflammation phase to the proliferation phase (4–10 days after the injury) (Figure 4). Treatment with Rg1 also induced protein expression of VEGF and TGF-β1 during the wound-healing inflammation phase (4–7 days after the injury), resulting in downregulation of expression at day 10 after the wound was fully healed (Figure 5). Other studies have shown that treatment with Rg1 increases angiogenesis and insulin secretion by increasing VEGF expression in the proliferative phase and progressively decreasing VEGF expression over time [38, 39]. In particular, Rg1 treatment reduced wound size compared to the control group (red ginseng extract, Rg1, and control mice were all diabetic). Factors such as TGF-β, PDGF, and interleukin (IL)-1 produced by T cells and macrophages stimulate a variety of growth factors; the expression of VEGF during the inflammatory stage also plays an important role in wound healing [40, 41, 42]. VEGF is an angiogenic cytokine that contributes to the proliferation phase of wound healing by promoting angiogenesis and the formation of fibrous cells, collagen, and granulation tissue around the wound [43, 44]. Many studies have focused on VEGF as a factor promoting angiogenesis and its role in various diseases [45, 46]. In the remodeling phase, VEGF induces cell degranulation, fibroblast proliferation, and migration of existing blood vessels by proteases; fibroblasts are transformed into α-smooth muscle actin bundles similar to smooth muscle cells, resulting in contraction of the wound [47, 48]. Nogami *et al.* [49] reported that VEGF mRNA levels were reduced significantly in diabetic animal models compared to normal models; thus, regulatory defects in VEGF delayed wound healing. Our results also showed that both red ginseng extract and Rg1 were affected by a delayed wound healing due to a lack of VEGF and TGF-β1 expression in untreated diabetic wounds (i.e., in the control group). In our histological analyses, both red ginseng and Rg1 treatment promoted formation of granulation tissue at days 7–10 after injury; wound contraction, reduction of inflammatory cells, and promotion of growth factors were observed during reconstruction at day 10. Treatment of tissues with Rg1 contributes to wound healing by promoting collagen production and VEGF expression [50, 51]. Red ginseng positively affects wound healing by modulating VEGF and TGF-β1 expression in diabetic wounds. Similarly, our results suggest that treatment with red ginseng may promote diabetic wound healing. In particular, Rg1 by regulates the expression of VEGF and TGF-β1; growth factors were also expressed at the inflammatory stage, promoting proliferation (Figure 3). This is the first report to show that red ginseng extract and Rg1 regulate the expression of VEGF and TGF-β1 to induce healing of diabetic wounds. Expression levels of these factors were confirmed by histological staining, IHC, and molecular biology.

## Conclusion

Rg1 treatment increases the expression of VEGF and TGF-β1, which are important for wound healing in STZ-induced diabetic wounds. Rg1 can improve diabetic wound healing by stimulating the production or activity of factors related to wound healing.

## Limitations

Although factors such as STZ can cause T1D, T2D is a complex metabolic disorder due to insulin resistance driven by environmental factors (e.g., obesity, and age). Additionally, animal experiments may not reflect all clinically observed complexities associated with disease [52]. Further studies of T2D may provide clinical data to corroborate the results obtained in this study.

## Acknowledgments

This research was supported by the Basic Science Research Program through the National Research Foundation of Korea (NRF) funded by the Ministry of Education (grant number 2019-061364) and the Soonchunhyang University research fund.

## Availability of data and materials

All study data used to support our findings are available from the corresponding authors upon request.

## Ethics approval and consent to participate

All procedures involving animals were in accordance with the ethical standards of the institution. The experimental design was approved by the Institutional Animal Care and Use Committee of Soonchunhyang University Medical School (SCHBC-Animal-2016–03).

